# A short, robust brain activation control task optimised for pharmacological fMRI studies

**DOI:** 10.1101/233783

**Authors:** Jessica-Lily Harvey, Lysia Demetriou, John McGonigle, Matthew B Wall

## Abstract

Functional magnetic resonance imaging (fMRI) is a popular method for examining pharmacological effects on the brain; however the BOLD response is an indirect measure of neural activity, and as such is vulnerable to confounding effects of pharmacological probes. Controlling for such non-specific effects in pharmacological fMRI studies is therefore an important consideration. We have developed two variants of a standardized control task that are short (5 minutes duration) simple (for both the subject and experimenter), widely applicable, and yield a number of readouts in a spatially diverse set of brain networks. The tasks consist of four functionally discreet three-second trial types (plus additional null trials) and contain visual, auditory, motor and cognitive (eye-movements, and working memory tasks in the two task variants) stimuli. Performance of the tasks was assessed in a group of 15 subjects scanned on two separate occasions, with test-retest reliability explicitly assessed using intra-class correlation coefficients. Both tasks produced robust patterns of brain activation in the expected brain regions, and reliability coefficients for the tasks were generally high, with four out of eight task conditions rated as ‘excellent’, and only one out of eight rated as ‘poor’. Voxel-wise reliability measures also showed good spatial concordances with the brain activation results. Either of the two task variants would be suitable for use as a control task in future pharmacological fMRI studies or for any situation where a short, reliable, basic task paradigm is required. Stimulus code is available online for re-use by the scientific community.

## Introduction

Functional Magnetic Resonance Imaging (fMRI) is currently one of the major standard methods in cognitive neuroscience research. FMRI provides reasonably high spatial and temporal resolution data, is flexible enough to accommodate a wide variety of experimental designs, and exposure to magnetic fields presents no danger to most subjects (Logothetis, 2008; Soares *et al*, 2016). FMRI can also be used as an index of pharmacological effects; drugs or hormones can be administered before or during a scanning session, and the results compared with a baseline or placebo session (e.g. Carhart-Harris *et al*, 2014; Comninos *et al*, 2017; Kaelen *et al*, 2016; Upadhyay *et al*, 2011). Pharmacological-fMRI studies may be used in the drug discovery process (Matthews *et al*, 2011; Wise and Tracey, 2006), in the characterization of the effects of commonly-prescribed drugs (Maron *et al*, 2016), or in the exploration of disorders such as addiction (Quelch *et al*, 2017).

Conducting pharmacological-fMRI investigations presents many of the same challenges as standard fMRI, but also has some unique issues. One fundamental concern is related to the fact that (most commonly) fMRI studies use the BOLD (Blood-Oxygen-Level-Dependent) signal as the primary end-point. This is a contrast produced by local changes in the ratio of oxygenated and de-oxygenated hemoglobin (Buxton *et al*, 1998; Friston *et al*, 2000), and is usually regarded as a proxy measure of neural activity. However, the relationship between neural activity and this vascular response (neurovascular coupling) is complex and relies on a number of cellular and metabolic processes (Logothetis *et al*, 2001). Use of a pharmacological agent combined with fMRI means that any differences observed in the BOLD response may be a combination of direct neural effects of the drug (usually the effects of interest), and indirect effects of the drug (e.g. on neurovascular coupling, or global, systemic effects on blood-pressure, cerebral blood flow, heart-rate, etc.; usually regarded as confounding effects). One example is caffeine which has direct neural effects on adenosine A_1_ and A_2a_ receptors, but is also a powerful cerebral vasoconstrictor (Diukova *et al*, 2012). Separating the neural and vascular effects of even such a selective and widely-studied drug as caffeine is therefore a considerable challenge. For detailed reviews of these issues see Bourke and Wall (2015), and Iannetti and Wise (2007).

One method of mitigating this problem is the use of an independent control task paradigm as part of a pharmacological fMRI scanning session (Iannetti and Wise, 2007). For example, Murphy *et al* (2009) used a visual control task in their study of the effect of citalopram on amygdala responses to emotional faces. In this case the lack of effect of the drug on the visual control task suggests that the effects seen in the main task are unlikely to be due to effects on neurovascular coupling, or other global/systemic effects. However, the use of a single (visual) control task, which gives activation in a subscribed region of the brain (the occipital lobe) is suboptimal as indirect effects on neurovascular coupling may still vary across the brain. Comninos *et al* (2017) used a much more elaborate control task (based on Pinel *et al*, 2007) in their recent study on the sex hormone kisspeptin. This task involved ten trial conditions which gave results in five separate functional domains (visual, auditory, language, motor, and cognitive), and in a much wider spatial distribution across the brain. This task involved relatively complex instructions for the subjects, and also included some culturally-specific language stimuli, which somewhat limits its broad applicability.

An ideal task for the control of pharmacological fMRI studies should have the following characteristics. First, it should be short in duration as it generally has to be included as part of a broader set of functional task paradigms, anatomical scans, and perhaps other MRI measures (resting-state fMRI, perfusion measures, spectroscopy etc.). Second, it should be simple, both for the subject to perform and for the experimenter to run and analyse. It should require no complex instructions, and depend upon only standard equipment (standard computer hardware/software, audiovisual systems, and simple response devices). Third, it should contain a number of different trial types, which produce activation in different brain networks, in as wide a spatial distribution across the brain as possible. This helps to rule out effects on neurovascular coupling which may differ in spatially remote brain regions. Fourth, it should be general-purpose; applicable to a wide range of different pharmacological fMRI studies. Fifth, it should be reliable; it should produce robust results within a single-session, and produce reliable results across multiple sessions. This last point is of particular importance, as use of an unreliable control task would constitute an additional confound, however no previous pharmacological fMRI study has explicitly assessed the reliability of its control task. Indeed reliability is relatively seldom formally assessed in fMRI studies (Plichta *et al*, 2012).

We have developed two variants of a task paradigm that meet the above mentioned criteria, and are furthermore programmed in an open-source software environment (PsychoPy; Peirce, 2007, 2008). One variant consists of visual, auditory, motor, and eye-movement trials. The other substitutes a brief working-memory task for the eye-movement trials, but is otherwise identical. Both are short (5 minutes in duration), simple (requiring only standard audiovisual equipment, and a single-button response box), and both produce four robust, distinct, and specific patterns of brain activation in widely-distributed brain regions. The reliability of the task variants across two scanning sessions has been explicitly assessed using a combination of voxel-wise and Region of Interest (ROI) based approaches.

## Methods

### Subjects

15 healthy subjects (6 males, 9 females) from ages 21-48 (mean age = 30) were scanned on two separate occasions with the average re-test interval being two weeks. All participants were fully briefed and provided informed consent.

### Task Design and Procedure

The tasks were programmed in PsychoPy (Peirce, 2007, 2008); a free, open-source, cross-platform Python library optimized for experimental design. The task consisted of 5 discreet trial types: auditory, visual, motor, cognitive and null trials, each lasting exactly three seconds. A small red, square fixation point was present throughout each task (except in one trial type, as noted below) at the centre of the screen. Auditory trials presented six pure tones for 0.5s each, at frequencies of 261.63Hz, 293.66Hz, 329.63Hz, 349.23Hz, 440Hz, and 493.88Hz (corresponding to the musical pitches C_4_, D_4_, E_4_, F_4_, A_4_, and B_4_, respectively). The order of the six tones was randomly determined on each trial. Visual trials consisted of a centrally-presented sine-wave grating subtending approximately 10° of visual angle and with a spatial frequency of 1.2 cycles/degree. The grating drifted laterally at a rate of 6 cycles per second, and the direction of drift reversed every 0.5s. Motor trials consisted of three presentations of a small image of a button, presented just above the centre of the screen, for 1s each. This was a cue for subjects to press the response box key, and the button image disappeared after each response was made. The ‘cognitive’ trial differed in the two variations of the task. In the eye-movement variant, the fixation point moved to six different locations corresponding to the compass locations North-East, East, South-East, North-West, West, and SouthWest. These points were mapped on a circle with a radius of approximately 8.75° of visual angle. Each location was maintained for 0.5s, and all six were presented (in a random order) in each three second trial. In the working-memory variant of the experiment, the cognitive trial consisted of a brief working memory task. This involved the presentation of two letter strings (containing four letters each), followed by a single letter. The subject’s task was to indicate whether the final, single letter was present in the first letter string. If the final letter was present in the first letter string, they were instructed to push the response button. If the final letter was not present in the first letter string they were instructed to make no response. For half the working memory trials the final letter was present in the first string, and for half it was not present. Finally, in the null trials the fixation point was maintained for three seconds, with no other stimuli presented.

The two task variants were identical, except for the inclusion of eye-movement trials in one, and working-memory trials in the other. Each task consisted of 100 trials (20 of each of the four active conditions, plus 20 null trials) presented in a standardized pseudo-random order. Separate versions of the two tasks reversed the trial order, and the order of presentation of these versions was counter-balanced across subjects and scans. The order of presentation of the two task variants in the scan sessions was also systematically varied across subjects and scans. The task durations were exactly five minutes (100 trials of 3s duration) plus a 10 second buffer period at the end.

Prior to each scan session, subjects were shown a demonstration version of each variant of the task, and instructed how to perform them. During the scanning session, visual stimuli were projected through a wave guide in the rear wall of the scanner room onto a screen mounted in the rear of the scanner bore. This was viewed in a mirror mounted to the head coil. Participants received auditory stimuli and instructions via MRI-compatible headphones, and responded using a one-button response box held in their right hand. Responses were recorded using PsychoPy’s data-logging routines.

### MRI data acquisition and analysis

Data were acquired on a Siemens 3T Magnetom Trio MRI scanner (Siemens Healthcare, Erlangen, Germany), equipped with a 32-channel phased-array head coil. A high-resolution T1-weighted image was acquired at the beginning of each scan using a magnetization prepared rapid gradient echo (MPRAGE) sequence with parameters from the Alzheimer’s Disease Research Network (ADNI; 160 slices × 240 × 256, TR = 2300 ms, TE = 2.98 ms, flip angle = 9°, 1 mm isotropic voxels, bandwidth = 240Hz/pixel, parallel imaging factor = 2; Jack *et al*, 2008). Functional data collection used an echo-planar imaging (EPI) sequence for BOLD contrast with 36 axial slices, aligned with the AC-PC axis (TR = 2000ms, TE = 31ms flip angle = 80°, 3mm isotropic voxels, parallel imaging factor = 2, bandwidth = 2298Hz/pixel). Each functional scan lasted five minutes and ten seconds and consisted of 155 volumes.

Analysis was completed with FSL version 5.0.4 (FMRIB’s software Library; Oxford Centre for Functional Magnetic Resonance Imaging of the Brain; www.fmrib.ox.ac.uk/fsl/). Anatomical Images were initially skull-stripped using BET (Brain Extraction Tool; included in FSL). Images were pre-processed with standard parameters (head-motion correction, 100 s temporal filtering, 6 mm spatial smoothing, co-registration to a standard template). First-level analysis used a General Linear Model (GLM) approach with the four active conditions modelled as separate regressors and the null trials implicitly modelled as the baseline. Also included were the first temporal derivatives of each time-series and head-motion parameters as regressors of no interest. Group level analyses computed a simple mean across all subjects and both scan sessions using FSL’s FLAME-1 model and a statistical threshold of *Z*=3.1, *p*<0.05 (cluster-corrected). Contrasts were defined to isolate the response to each trial type relative to the null trials (baseline sections of the time-series). Two separate sets of analyses were conducted, for data from the two task variants.

Additional analyses used Intra-Class Correlation (ICC) coefficients to assess the reliability of responses across the two scanning sessions. This was performed in two ways; using an ROI-based approach, and by generating statistical maps of ICC values in a voxel-wise manner. For the ROI analysis, five regions were defined based on expected locations of brain activation in the tasks: primary auditory cortex in the superior temporal lobe (bilateral; auditory trials), primary visual cortex in the calcarine sulcus (bilateral; visual trials), left-hemisphere motor cortex (motor trials), the Frontal Eye Fields (FEF; bilateral; eye-movement trials), and the Dorso-Lateral Pre-Frontal Cortex (DLPFC; bilateral; working memory trials). ROIs were defined as 5mm-radius spheres, and positioning coordinates were determined using guidance from relevant meta-analytic terms on Neurosynth (http://neurosynth.org/). The ROI definition was therefore performed completely independently from the main experimental data. Activation amplitude data was extracted from these ROIs for all subjects/scans and ICC(3,1) statistics were calculated using SPSS (IBM Corp; Armonk, NY).

The ICC statistical maps were produced using custom Python code and produced voxelwise images of ICC(3,1) statistics. For the purposes of thresholding the results, the ICC values were then transformed into standardized values (Z scores) using the method of Fisher (1915). These images were then thresholded using the same statistical criterion used for the group level BOLD activation analyses; *Z* > 3.1, *p* < 0.05 (cluster-corrected for multiple comparisons). These thresholded images were then used to mask the original ICC voxelwise images, to finally produce a robustly thresholded image, which also retains the original, more intuitive, ICC values.

## Results

### Behavioural performance

Subjects’ behaviour was recorded and analysed to verify compliance with the task demands. An average accuracy rate of 93% was achieved within the working memory task. 94% and 97% accuracy was achieved for the motor task within the eye movement variant and working memory variant, respectively. All subjects performed the tasks satisfactorily.

### Group-level task activation

All tasks performed as expected and produced robust patterns of brain activity in regions previously shown to be activated by similar tasks. Performance of auditory, visual, and motor components of the tasks was consistent across both task variants (see figures 1a and 2a). Auditory trials produced strong bilateral activation within the superior temporal regions, consistent with primary auditory cortex (Robson *et al*, 1998). Visual trials produced activity in posterior calcarine sulcus and the occipital pole (primary visual cortex), and in the lateral visual region V5/MT+ (Smith *et al*, 2006; Wall *et al*, 2008). Motor trials produced activity in the left-hemisphere post-central sulcus, consistent with the known location of the hand representation in primary motor cortex (Lotze *et al*, 2000).

In the eye-movement variant of the experiment, the eye-movement task produced activation in the Frontal Eye Fields (FEF), alongside activity within V5/MT+, the anterior portion of the calcarine sulcus/primary visual regions, and the intraparietal sulcus (see figure 1). This is generally consistent with previous reports of brain activity associated with eye-movement tasks. In the working-memory variant of the experiment, the working memory trials produced a highly robust activation pattern corresponding closely to that shown in conventional working memory tasks, such as the N-back (Owen *et al*, 2005). These regions included bilateral DLPFC, intraparietal sulcus, superior parietal lobule, dorsal anterior cingulate and the temporo-parietal junction (see figure 2).

**Figure 1.**
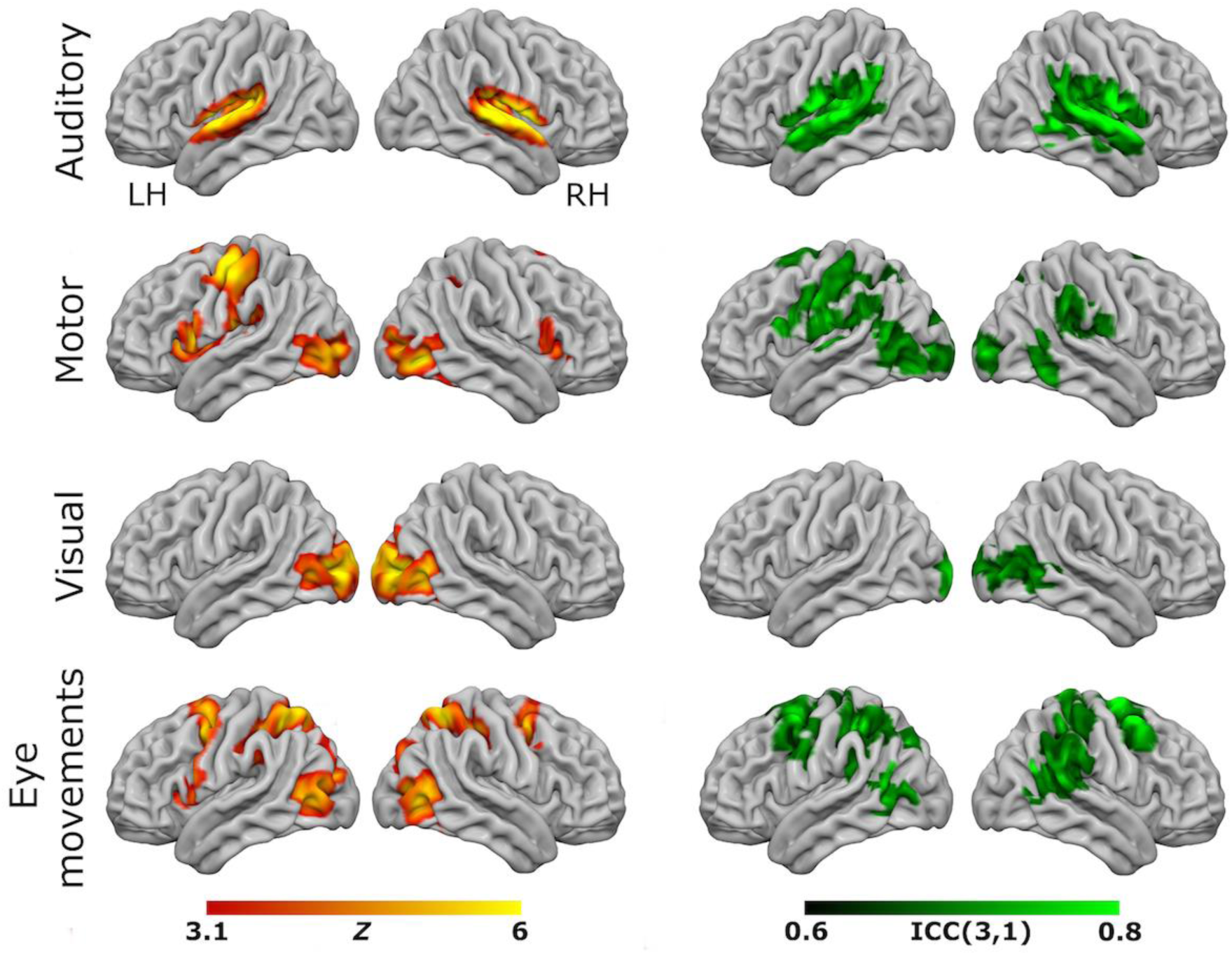
Results from the eye-movement variant of the task paradigm. Results of group-level analyses represented on a cortical surface rendering of a standard anatomical image (MNI152). Left column: Active brain regions for each contrast (mean of both scanning sessions) with functional maps thresholded at *Z* > 3.1, *p* < 0.05 (cluster-corrected). Right column: Results of the reliability analysis comparing session 1 to session 2; Intra-class correlation (3,1) maps, masked with a *Z*-transformed, thresholded (*Z* > 3.1, *p* < 0.05; cluster-corrected) version in order to produce a robustly-thresholded image, while retaining the original ICC values (see methods for full details). Rows 1-4 are auditory, motor, visual and eye-movement trials.

**Figure 2.**
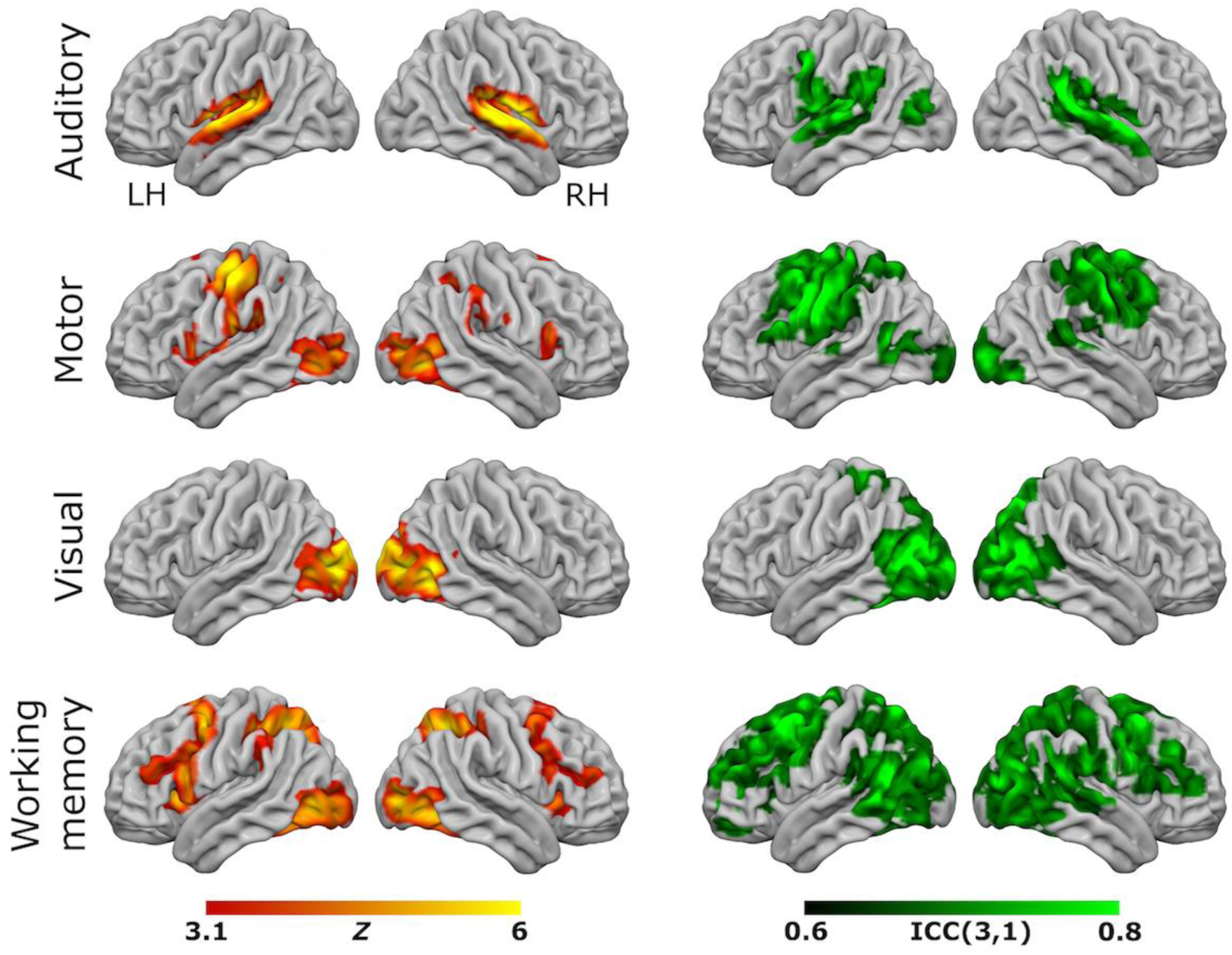
Results from the working-memory variant of the task paradigm. Results of group-level analyses represented on a cortical surface rendering of a standard anatomical image (MNI152). Left column: Active brain regions for each contrast (mean of both scanning sessions) with functional maps thresholded at *Z* > 3.1, *p* < 0.05 (cluster-corrected). Right column: Results of the reliability analysis comparing session 1 to session 2; Intra-class correlation (3,1) maps, masked with a *Z*-transformed, thresholded (*Z* > 3.1, *p* < 0.05; cluster-corrected) version in order to produce a robustly-thresholded image, while retaining the original ICC values (see methods for full details). Rows 1-4 are auditory, motor, visual and working memory trials.

Parameter estimate data were extracted from each contrast using a set of five ROIs: primary auditory cortex (auditory trials), frontal eye-fields (eye-movement trials), left-hemisphere primary motor cortex (motor trials), primary visual cortex (visual trials), and dorsolateral-prefrontal cortex (working memory trials). These data are plotted for each condition and scan session in figure 3. Statistical analysis of these data used paired t-tests to compare data from each contrast across the two scanning sessions, and a Bonferroni-corrected alpha value of *p* < 0.00625 (corrected for 8 comparisons). None of the comparisons showed significant results except for auditory trials, in the eye-movement variant; *t* (14) = 3.341, *p* = 0.00485.

**Figure 3.**
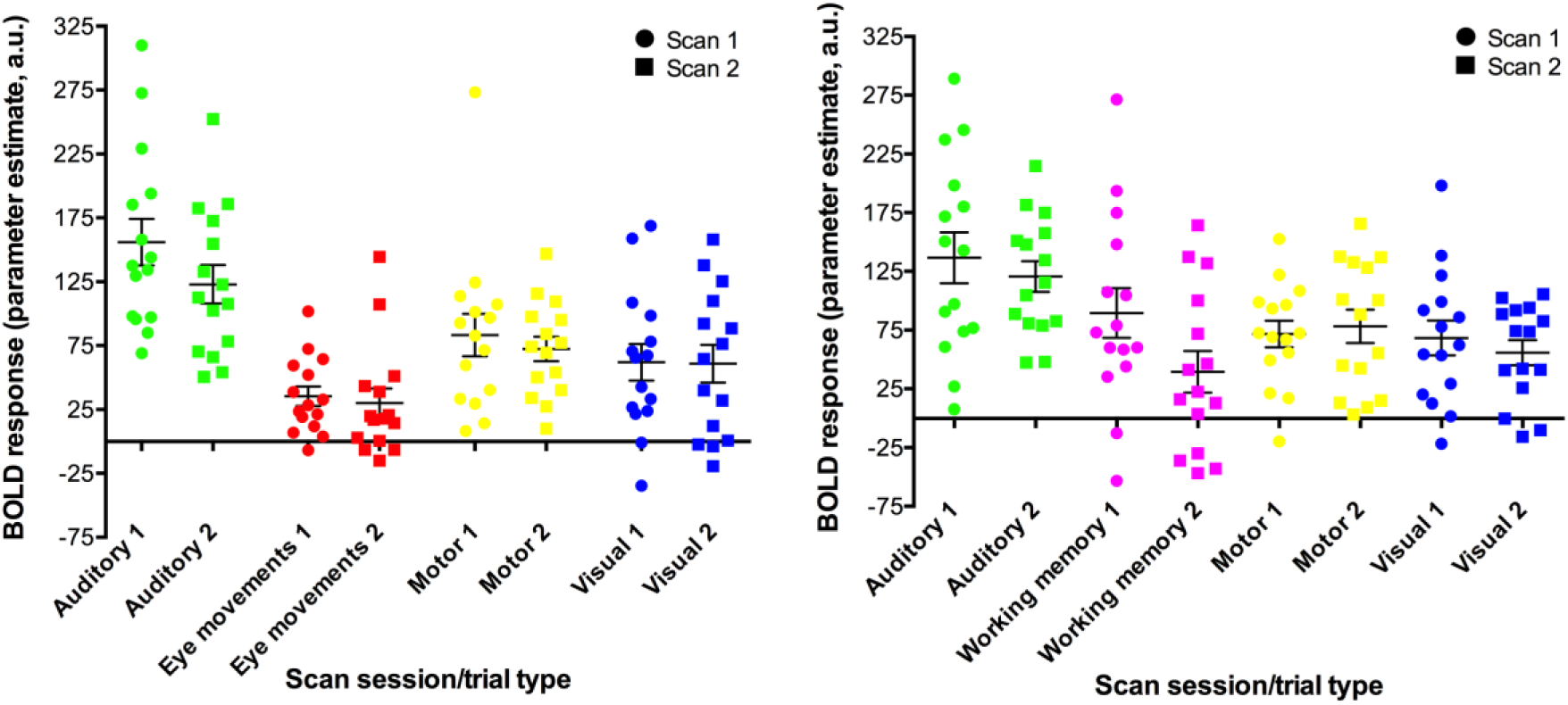
ROI data for each task condition within the two task variants (left panel = eye-movement variant, right panel = working-memory variant). Units are parameter estimates resulting from each of the four contrasts in each GLM analysis, relative to baseline (null trials) and are arbitrary units. ROIs are primary auditory cortex (auditory trials; green), frontal eye-fields (eye-movement trials; red), left-hemisphere primary motor cortex (motor trials; yellow), primary visual cortex (visual trials; blue), and dorsolateral-prefrontal cortex (working memory trials; pink). See supplementary figure 3 for images of the ROIs.

### Reliability analyses

To assess voxel level reliability, intra-class correlation (ICC(3,1)) maps were created for each task (figures 1 and 2; right columns). These show a spatial distribution very similar to the activation maps, with peak reliability estimates generally corresponding to the location of peak task-related activation. Reliability estimates in the working-memory variant of the task were generally higher and more widespread than in the eye-movement variant. For additional visualizations of the spatial correspondence between the activation maps and the ICC results, see supplementary figures 1 and 2.

In the ROI analysis, 4/8 ROIs featured ICC values of 0.75 or above, which is classed as ‘excellent’ under Cicchetti’s (1994) scheme for interpretation of ICC results. A further three ROIs had values in the range 0.4-0.59 which is classed as ‘fair’ reliability. Only one was < 0.4, and thus classed as ‘poor’. The auditory task featured the most robust reliability, with values of 0.849 in the eye-movement variant and 0.840 in the working-memory variant. The DLPFC ROI showed strong reliability of 0.589 for the working-memory task, and the FEF ROI had a similar score of 0.524 for the eye-movement task. Reliability within the primary visual cortex ROI was relatively low in the eye-movement variant of the task (0.466), however this ROI was highly reliable (0.765) in the working-memory variant. A similar dissociation was seen in the left motor cortex ROI with relatively poor reliability seen in the eye-movement variant (0.258) but much higher reliability (0.778) in the working-memory variant (Table 1).

Unthresholded statistical maps resulting from all the group-level analyses (brain activation maps, and the voxel-wise ICC maps) are available to view at: https://neurovault.org/collections/3264/

**Table 1.** ICC(3,1) values for the different trial conditions, in both variants of the experiment. Values in bold are classed as having ‘excellent’ reliability, those in italics are classed as having ‘fair’ reliability (Cicchetti, 1994).

| Eye-movement variant |  | Working memory variant |  |
| --- | --- | --- | --- |
| Task condition | ICC (3,1):<br>Scan 1 vs. Scan 2 | Task condition | ICC (3,1):<br>Scan 1 vs. Scan 2 |
| Auditory | **0.849** | Auditory | **0.84** |
| Visual | *0.434* | Visual | **0.765** |
| Motor | 0.258 | Motor | **0.778** |
| Eye-movement | *0.524* | Working memory | *0.589* |

## Discussion

We have developed and successfully validated two variants of a novel fMRI control task and demonstrated that they show high test-retest reliability. These tasks are short (five minutes duration), relatively simple for both the experimenter and subject (they require only standard audio-visual presentation equipment and a one-button response box), highly robust in terms of the amplitude of brain activation produced, and show strong reliability features across two sessions. Each variant also produces a number of useful readouts (visual, auditory, motor, cognitive/eye-movements) in a wide spatial distribution across the brain.

Both task variants performed similarly for visual, auditory, and motor trials, with robust activity seen in primary visual, auditory, and motor cortex respectively, and little ‘off-target’ activation evident. The eye-movement task also produced a characteristic pattern of brain activity similar to that seen in previous eye-movement studies (e.g. Berman *et al*, 1999). The working memory task, though only requiring a very brief (two-second) retention interval, produced a highly similar pattern of activity to that seen in more standard working memory tasks such as the N-back task (Owen *et al*, 2005).

Importantly, reliability of the tasks was also assessed, and found to be generally high. Reliability assessment using ICC (or other measures) is still relatively uncommon for fMRI experiments, but is an important step in validating task paradigms (Caceres *et al*, 2009). The ICC measures obtained here compare very favourably with previous reports using auditory and working memory tasks (Caceres *et al*, 2009), a cognitive-emotive test battery (Plichta *et al*, 2012), and a reward task (Fliessbach *et al*, 2010). However, some task conditions were seen to be more reliable than others. In particular, reliability in the working-memory variant of the experiment was generally higher than in the eye-movement variant. One possible explanation for this difference may be due to the much more cognitively demanding features of the working-memory variant, which led to a higher level of attention and engagement to all the task conditions in that variant. The high reliability, short duration, and ease of use of these tasks make them ideal for inclusion as control tasks in pharmacological-MRI studies, as suggested by Iannetti and Wise (2007), and Bourke and Wall (2015). Inclusion of tasks which are (hypothetically) unaffected by the drug helps rule out alternative explanations related to systemic drug effects (on blood pressure, heart-rate, etc.), effects on local vasculature, or neurovascular coupling; all of which can theoretically modulate the BOLD response. One previous study investigating modulation of amygdala responses by citalopram (Murphy *et al*, 2009) used a simple checker-board visual control task. Use of a single control task where activation is restricted to the occipital lobe is sub-optimal as the drug may potentially still produce non-neural effects in other brain regions. A recent study on the brain effects of the sex hormone kisspeptin (Comninos *et al*, 2017) used a control task with a number of readouts in different brain regions (based on Pinel *et al*, 2007). This task was complex, with ten individual stimulus conditions, different response options, and contained high-level cognitive stimuli (performing mental arithmetic, reading sentences on the screen, and listening to recorded voices) which included culture- and language-specific features. This complexity and the use of language-specific stimuli limit the broad applicability of this task.

The tasks evaluated here represent a good compromise between ease of use, wide applicability, a short duration, reliable results, and the desirability of providing a number of readouts in spatially diverse brain regions. While the working memory variant appears to be somewhat more robust, more reliable, and produces a wider pattern of brain activity, it is also more cognitively demanding and has significantly more complex instructions. This may make it less suitable for any patient group with significant cognitive impairments, who may struggle with a fast, demanding task. The eye-movement variant may therefore be more suitable for these groups. Additionally, the eye-movement variant may also be more suitable where the drug under investigation is hypothesized to have an effect on cognition. In this case, the working-memory variant may be inappropriate as a control task, as it strongly engages well-known cognitive brain regions. Either variant would also be suitable for use in a number of other situations where a short, reliable fMRI task that yields a number of readouts is required, for example in systematic testing of fMRI acquisition sequence parameters (as in Demetriou *et al*, 2016).

We have evaluated two variants of a novel task paradigm, suitable for use as a control task in pharmacological fMRI studies, or for any use where a general-purpose battery of basic tasks/stimuli is required. The tasks produce robust brain activation and have strongly favourable reliability features. The tasks are programmed in an open-source language and experimental presentation application (Python/PsychoPy), and we have therefore made the stimulus code freely available at https://figshare.com/articles/fMRIcontroltaskzip/5162065 (DOI: 10.6084/m9.figshare.5162065; Google-generated short-link: goo.gl/DAqn4V). We encourage any interested researchers to download the programs and use them in their research.

## Acknowledgements

We would like to thank the Imanova Center for Imaging Sciences (Hammersmith Hospital, London, UK) for the scanner time required to complete the project, and general support throughout the investigation

---

**Supplementary figure 1.**
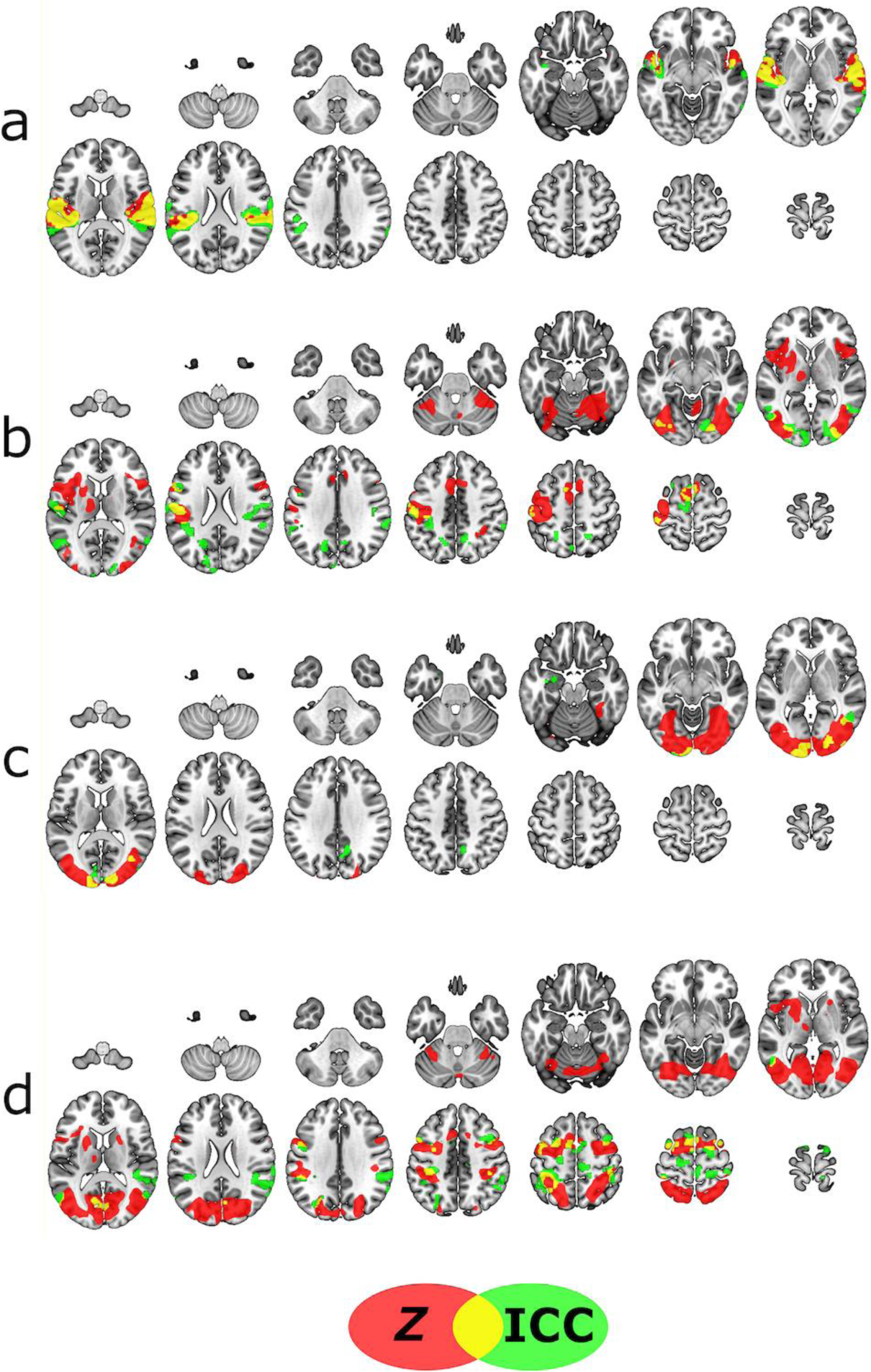
BOLD activation data (Z-scores) and ICC(3,1) reliability values (both thresholded at *Z* > 3.1, *p* < 0.05, cluster-corrected) from the eye-movement variant represented on the same anatomical image in order to visualize the spatial relationship between the two sets of data. a) Auditory trials. b) Motor trials. c) Visual trials. d) Eye movement trials.

**Supplementary figure 2.**
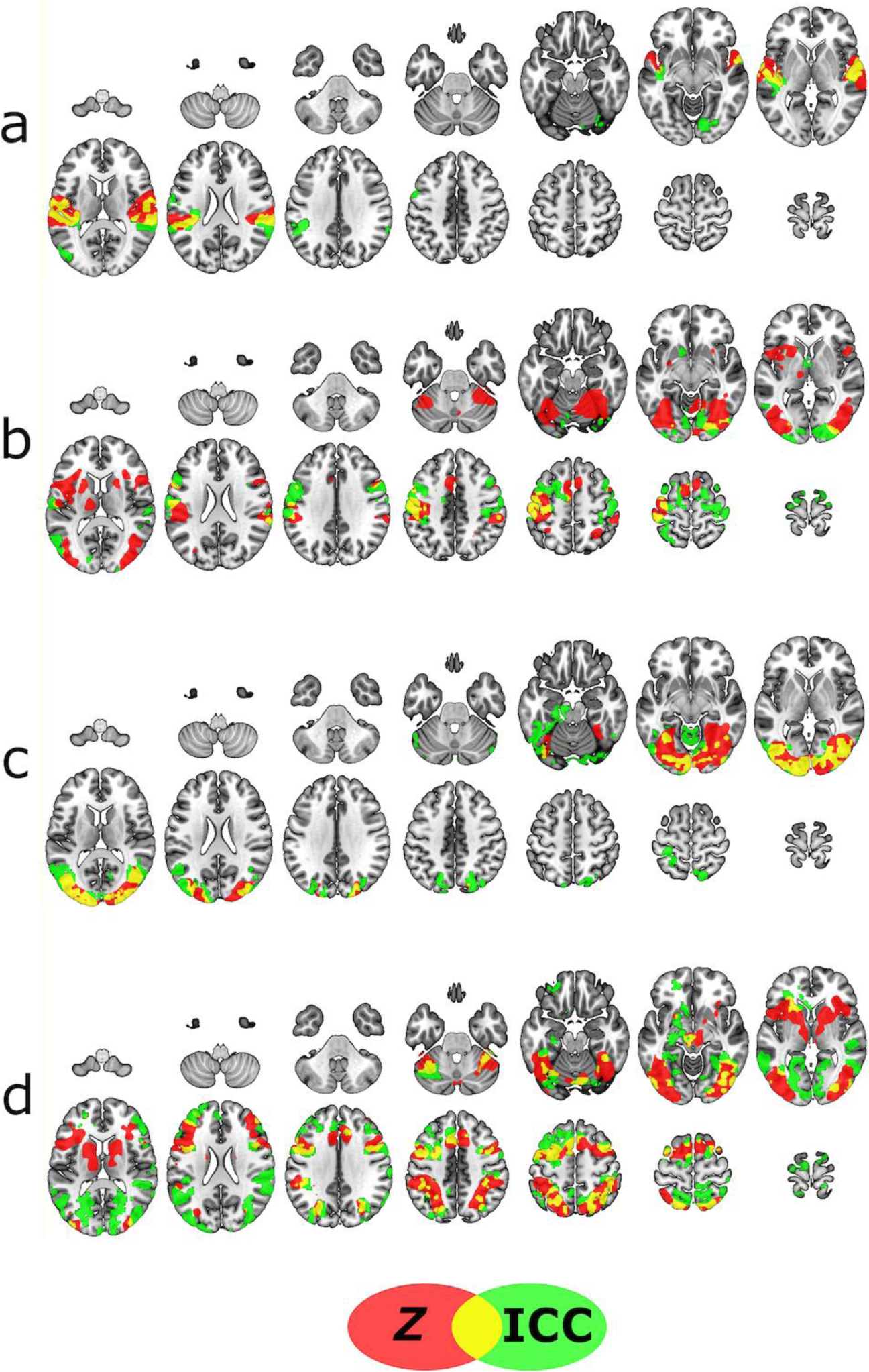
BOLD activation data (Z-scores) and ICC(3,1) reliability values (both thresholded at *Z* > 3.1, *p* < 0.05, cluster-corrected) from the working-memory variant represented on the same anatomical image in order to visualize the spatial relationship between the two sets of data. a) Auditory trials. b) Motor trials. c) Visual trials. d) Working memory trials.

**Supplementary figure 3.**
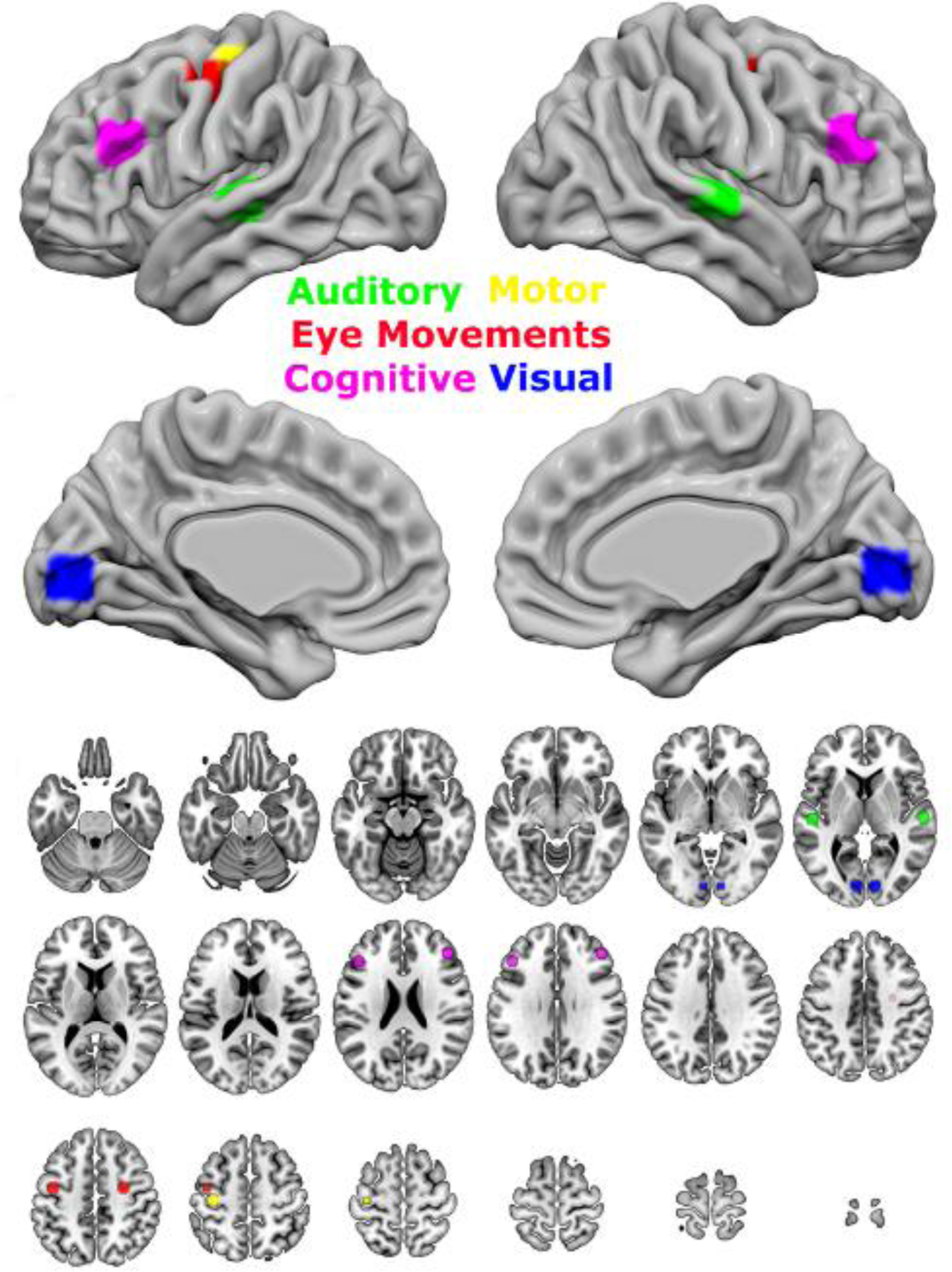
Regions used in the ROI analysis visualized on the cortical surface (upper panel) and on a set of axial slices (lower panel). ROIs were independently defined as 5mm-radius spheres, using positioning coordinates determined using guidance from relevant meta-analytic terms on Neurosynth (http://neurosynth.org/).

